# Exploring miRNAs as the key to understand symptoms induced by ZIKA virus infection through a collaborative database

**DOI:** 10.1101/042382

**Authors:** Victor Satler Pylro, Francislon Silva de Oliveira, Daniel Kumazawa Morais, Sara Cuadros Orellana, Fabiano Sviatopolk-Mirsky Pais, Julliane Dutra Medeiros, Juliana Assis Geraldo, Jack Gilbert, Angela Volpini, Gabriel da Rocha Fernandes

**Author notes:** These authors contributed equally to this work (Victor Satler Pylro, Francislon Silva de Oliveira).

## Abstract

Zika virus (ZIKV) is an emerging mosquito-borne flavivirus, first isolated in 1947 from the serum of a pyrexial rhesus monkey caged in the Zika Forest (Uganda/Africa)^1^. In 2007 ZIKV was reported to of been responsible for an outbreak of relatively mild disease, characterized by rash, arthralgia, and conjunctivitis on Yap Island, in the western Pacific Ocean^2^. In the past year, ZIKV has been circulating in the Americas, probably introduced through Easter Island (Chile), by French Polynesians^3^. In early 2015, a new outbreak was recognized in northeast Brazil^4^, where concerns over its possible links with infant microcephaly have been discussed^5,6^. Providing a definitive link between ZIKV infection and birth defects is still a big challenge^7^. MicroRNAs (miRNAs), are small noncoding RNAs that regulating post-transcriptional gene expression by translational repression, and play important roles in viral pathogenesis^8^ and brain development^9^. It is estimated that more than 60% of human protein-coding genes contain at least one conserved miRNA-binding site^10^. The potential for flavivirus-mediated miRNA signaling dysfunction in brain-tissue develop provides a compelling mechanism underlying perceived linked between ZIKV and microcephaly. Here, we provide strong evidences toward to understand the mechanism in which miRNAs can be linked to the “congenital Zika syndrome” symptoms. Moreover, following World Health Organization (WHO) recommendations^11^, we have assembled a database that could help target mechanistic investigations of this possible relationship between ZIKV symptoms and miRNA mediated human gene expression, helping to foster potential targets for therapy.

---

Two hypotheses of how miRNAs can influence ZIKV/human-host interaction were raised to support our database. First, viruses can transcribe miRNAs that enable them to benefit from cellular and viral gene expression (e.g. Herpesvirus, Polyomavirus, Ascovirus, Baculovirus, Iridovirus, Adenovirus families)^12,13^. RNA retrovirus miRNAs are encoded through a RNA polymerase III (pol III) promoter, instead of being transcribed by pol II. Virus-encoded miRNAs can play a role in persistent infections through subtle modulation of gene expression, leading to prevention of host cell death, evasion of the host immune system and regulation of the latent-lytic switch^14^. Second, retrovirus genomes can directly interact with cellular miRNAs to enhance viral replication potential through a mechanism of innate antiviral immunity^12^. In this sense, by recruiting/exploiting cellular miRNAs, an RNA virus can disturb normal host gene expression regulation, which can trigger different molecular diseases. To enable testing of these hypotheses, the *ZIKV collaborative database* (ZIKV-CDB) was assembled to allow for: (i) searching for predicted ZIKV miRNAs mimicking human miRNAs [searching criteria includes: “Gene name”, “Gene Symbol” or “Ensembl ID”] (hypothesis 1); and (ii) searching for human miRNAs with possible binding-sites to the ZIKV genomes (hypothesis 2).

The ZIKV-CDB comprises miRNAs predicted using HHMMiR^15^ for all complete ZIKV genomes currently available at the GenBank (February, 2016 - http://www.ncbi.nlm.nih.gov). Hairpin prediction was also performed for all *de novo* miRNAs using previously predicted RNA secondary structure^16^, and mature miRNAs were delineated with PHDcleav^17^. Finally, to test hypothesis 1, potential human genome (Ensembl GRCh37) target sites for the predicted ZIKV miRNAs were detected with miRanda^18^ using default parameters (minimum score=140; minimum energy=1). To test hypothesis 2, we retrieved all mature human miRNA sequences from miRBase Sequence Database (Release 21 – http://www.mirbase.org), and mapped these against the available ZIKV genomes using miRanda^18^ with default parameters, to keep only those miRNAs with a minimum complementarity to ZIKV genomes [at least with complementarity to the miRNA seed region (6-10 nt) of the miRNA]. The ZIKV-CDB can be accessed through a web interface at http://zikadb.cpqrr.fiocruz.br.

The ZIKV-CDB has enabled us to predict miRNAs from ZIKV that have high-confidence targets in the human genome (Table 1), as well as cellular (human) miRNAs able to interact with ZIKV genomes (Table 2). This knowledge base should help to delineate targets that have the potential to affect the expression of genes associated with microcephaly and other neurodevelopmental syndromes. Some of these targets include, the peroxisomal biogenesis factor 26 gene (PEX26), the fibroblast growth factor 2 (FGF2), the SET binding factor 1 (SBF1), the hook microtubule-tethering protein 3 (Hook3) and the pleckstrin homology and RhoGEF domain containing G4 (PLEKHG4) (Table 1). PLEKHG4 polymorphisms have been related to spinocerebellar ataxia^19^, a progressive-degenerative genetic disease. Also, Hook3 has been reported to interact with Pericentriolar Material 1 (PCM1) during brain development, and an imbalance in the Hook3-PCM1 interaction can cause premature depletion of the neural progenitor pool in the developing neocortex^20^. Finally, defects in the PEX26 gene can lead to a failure of protein import into the peroxisomal membrane or matrix, being the cause of several neuronal disorders, including Zellweger syndrome (ZWS), and neonatal adrenoleukodystrophy (NALD)^21^, ^22^.

**Table 1.**
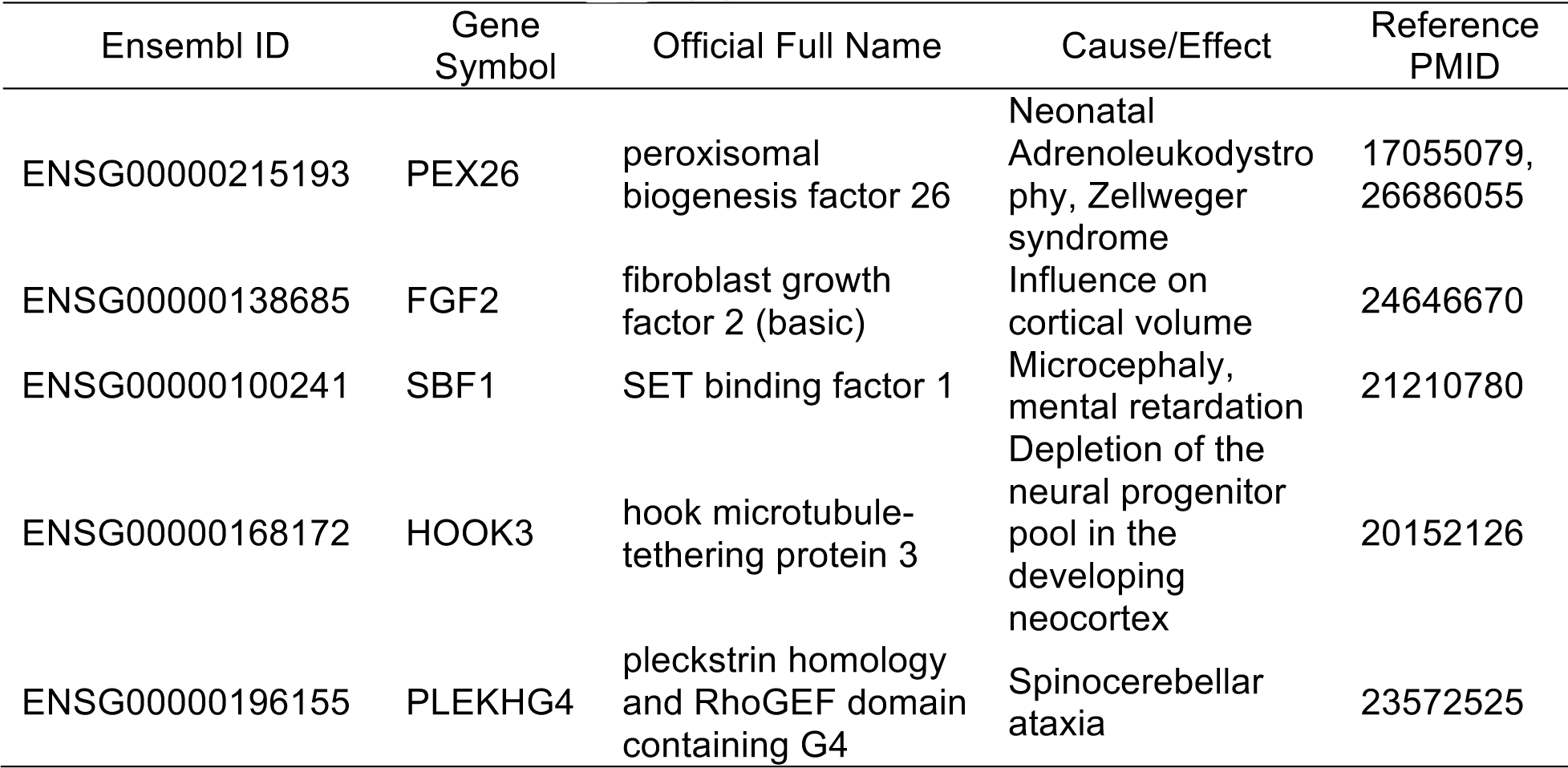
Examples of human targeting-genes to ZIKV predicted miRNAs.

| Ensembl ID | Gene Symbol | Official Full Name | Cause/Effect | Reference PMID |
| --- | --- | --- | --- | --- |
| ENSG00000215193 | PEX26 | peroxisomal biogenesis factor 26 | Neonatal Adrenoleukodystrophy, Zellweger syndrome | 17055079, 26686055 |
| ENSG00000138685 | FGF2 | fibroblast growth factor 2 (basic) | Influence on cortical volume | 24646670 |
| ENSG00000100241 | SBF1 | SET binding factor 1 | Microcephaly, mental retardation | 21210780 |
| ENSG00000168172 | HOOK3 | hook microtubule-tethering protein 3 | Depletion of the neural progenitor pool in the developing neocortex | 20152126 |
| ENSG00000196155 | PLEKHG4 | pleckstrin homology and RhoGEF domain containing G4 | Spinocerebellar ataxia | 23572525 |

**Table 2.** Examples of human miRNAs able to interact with ZIKV genomes.

| miRNAs | Description* | Reference PMID |
| --- | --- | --- |
| hsa-miR-34a | CDK6 target | 23918663 |
| hsa-mir-324-3p | Cdk5rap2/MCHP3 target | 21632253<br>23726037 |
| hsa-mir-15b-5p | MCPH1/BRIT1 target | 12046007 |
| hsa-mir-21-5p | MCPH4/CEP152 target | 20598275 |
| hsa-mir-335-5p | MCPH1/BRIT1 target | 12046007 |
| hsa-mir-615-3p | WDR62/MCPH2 target | 20890278 |
| hsa-miR-193b-3p | WDR62/MCPH2/ target | 20890278 |

\*All these genes have been previously related to microcephaly.

*All these genes have been previously related to microcephaly.

Interestingly, several human miRNAs known to exert an influence on the expression of genes with a functional role in neuronal development, were found to have sequence complementarity to regions in the ZIKV genomes (Table 2). The human hsa-miR-34a miRNA has, as one of its targets, the Cyclin-Dependent Kinase 6 (CDK6) gene, which is technically able to interact with the ZIKV genome. CDK6 is associated with the centrosome during mitosis, controlling the cell cycle division phases in neuron production^23^. Mutation in CDK6 can lead to a deficient centrosomes division, which in turns can cause autosomal recessive primary microcephaly (MCHP)^23^. There are seven well-know genes encoding centrosomal proteins that are involved in the autosomal recessive primary microcephaly (MCPH)^24^, including the CDK5 Regulatory Subunit Associated Protein 2 (Cdk5rap2 or MCHP3) gene. We found a possible binding-site to the hsa-mir-324-3p, a cellular miRNA targeting Cdk5rap2 gene, in the ZIKV genomes. Equally, we found that ZIKV genomic regions can bind the hsa-mir-615-3p and hsa-miR-193b-3p human miRNAs, which target the WD Repeat Domain 62 (WDR62 or MCHP2), also related to MCHP when mutated. Remarkably, a hsa-mir-21-5p miRNA complementary site was found on the genomes of ZIKV from Brazil, Haiti, Martinique and French Polynesia, but not in those from Africa. This miRNA targets the MCHP4 gene, also linked to microcephaly cases. This geographic and hence historical accumulation of genomic differences may explain the recent rise of microcephaly, and also corroborates the predicted pathway of transmission from Africa, through Oceania, and into Central and South America.

To further support this, phylogenetic analysis was performed for all complete ZIKV genomes (Figure 1), which identified a cluster of strains isolated in Americas and Oceania (“recent” strains), and another with the African strains (“ancient” strains). A third group, including other two Flavivirus’ close related to ZIKV, was added as out-group. These difference were also supported by phylogenetic analysis of only the predicted miRNAs encoded by each ZIKV strain. Nine predicted miRNA types were identified with types 1-4 restricted to recent strains and 5-8 restricted to ancient strains. Type 9 was only found in strains from Central Africa Republic (Figure 2). The predicted miRNAs could target 14,745 human genes; 9,106 were specific to the ‘ancient’ strains, 2,840 specific to the ‘recent’ strains, and 2,789 were shared.

**Figure 1.**
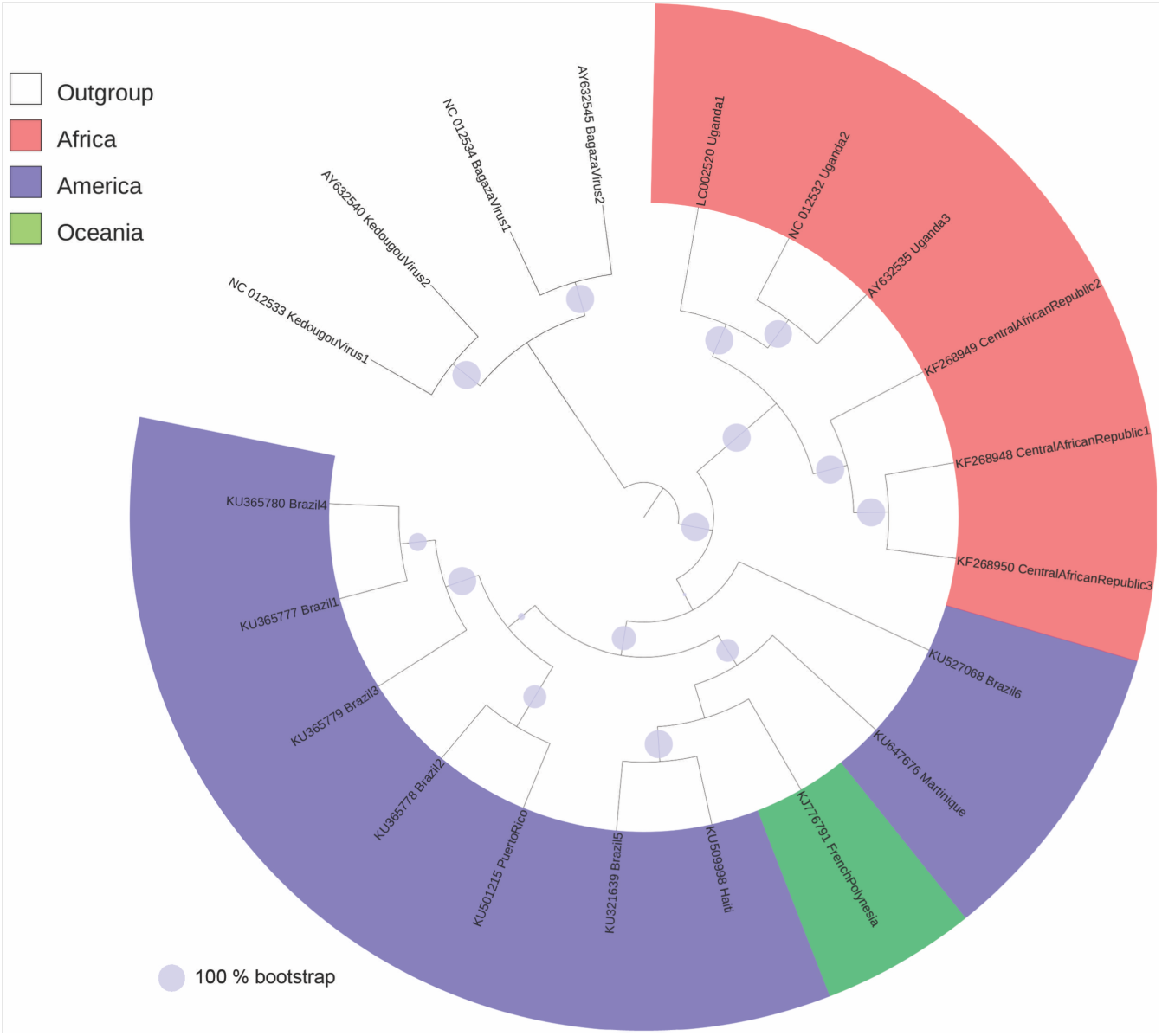
Phylogenetic analysis of all complete genomes of Zika Virus, available at GenBank (February, 2016). The GenBank accession number and the country of origin are indicated on the ZIKV branches for all strains, except for those from de out-group, where the name of the viruses is provided [Kedougou virus (NC_012533 and AY632540); Bagaza virus (AY632545 and NC_012534)]. The size of the full circles on the branches means percentage of bootstrap (100 replicates).

**Figure 2.**
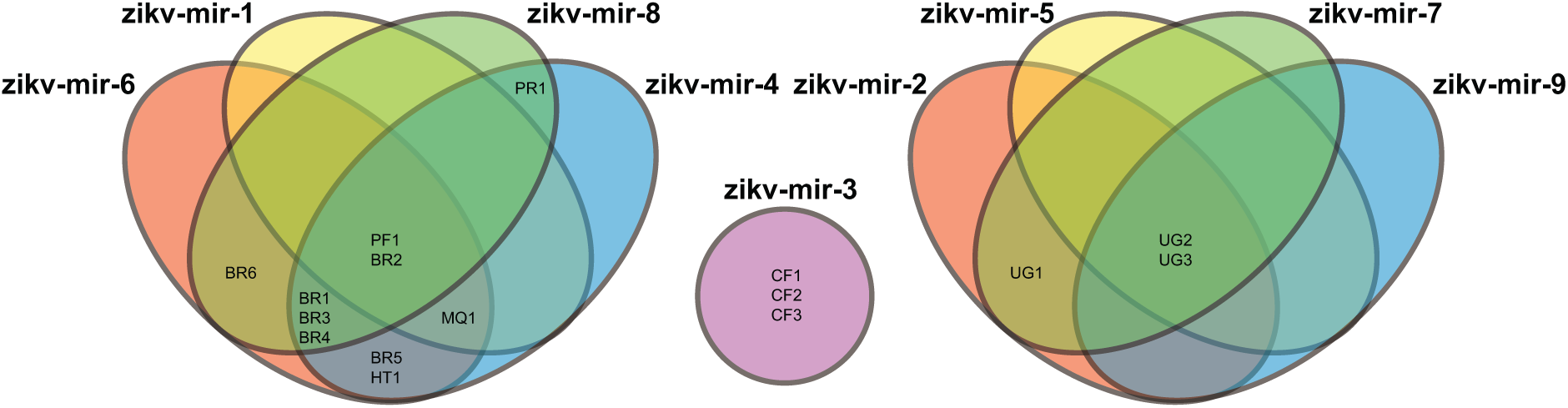
Venn diagram showing the nine different types of miRNAs predicted to be encoded by all analyzed ZIKV genomes. Brazil (BR); Central African Republic (CF); French Polinesia (FP); Haiti (HT); Martinique (MQ); Puerto Rico (PR); Uganda (UG).

Recently, ZIKV was isolated from the brain tissue of a fetus diagnosed with microcephaly^25^. However, the mechanism by which ZIKV alters neurophysiological development remains unknown, preventing therapeutic intervention. Our results provide new insights associated with the potential influence of miRNAs on the expression of human-genes associated with the symptoms of “congenital ZIKA syndrome”. The ZIKV-CDB provides a crucial knowledge base to mitigate the impacts of this emerging health problem^26^. ZIKV-CDB is an open-source and collaboration-based forum for sharing and identification of potential mechanistic and therapeutic targets. These data will guide experimental investigation to elucidate any association between ZIKV infection and neurobiological development in infants. The ZIKV-CDB will be constantly maintained and curated by the staff of the Genomics and Computational Biology Group, from FIOCRUZ/CPqRR (http://www.cpqrr.fiocruz.br).

